# pyVIPER: A fast and scalable Python package for rank-based enrichment analysis of single-cell RNASeq data

**DOI:** 10.1101/2024.08.25.609585

**Authors:** Alexander L.E. Wang, Zizhao Lin, Luca Zanella, Lukas Vlahos, Miquel Anglada Girotto, Aziz Zafar, Heeju Noh, Andrea Califano, Alessandro Vasciaveo

## Abstract

**Summary:** Single-cell sequencing has revolutionized biomedical research by offering insights into cellular heterogeneity at unprecedented resolution. Yet, the low signal-to-noise ratio, characteristic of single-cell RNA sequencing (scRNASeq), challenges quantitative analyses. We have shown that gene regulatory network (GRN) analysis can help overcome this obstacle and support mechanistic elucidation of cellular state determinants, for example by using the VIPER algorithm to identify Master Regulator (MR) proteins from gene expression data. A key challenge, as the size and complexity of scRNASeq datasets grow, is the need for highly scalable tools supporting the analysis of large-scale datasets with up to hundreds of thousands of cells. To address it, we introduce pyVIPER, a fast, memory-efficient, and highly scalable Python toolkit for assessing protein activity in large-scale scRNASeq datasets. pyVIPER supports multiple enrichment analysis algorithms, data transformation/postprocessing modules, a novel data structure for GRNs manipulation, and seamless integration with AnnData, Scanpy and several widely adopted machine learning libraries. Compared to VIPER, benchmarking reveals orders of magnitude runtime reduction for large datasets—i.e., from hours to minutes— thus supporting VIPER-based analysis of virtually any large-scale single-cell dataset, as well as integration with other Python-based tools.

**Availability and Implementation:** pyVIPER is available on GitHub (https://github.com/alevax/pyviper) and PyPI (https://pypi.org/project/viper-in-python/).

**Supplementary information:** Supplementary data are available at Bioinformatics online. Accompanying data for the tutorials are available on Zenodo (https://zenodo.org/records/10059791).

## 1 Introduction

We have shown that the analysis of gene expression data through the interrogation of context-specific gene regulatory networks (GRNs) plays a key role in the identification of Master Regulator (MRs) proteins representing mechanistic determinants of cellular states (Califano and Alvarez, 2017; Li, *et al*., 2023; Malagola, *et al*., 2024; Vasciaveo, *et al*., 2023). These MR proteins can be computationally identified using the VIPER (Virtual Inference of Protein Activity by Enriched Regulon analysis) and metaVIPER algorithms through the analysis of their transcriptional targets (Alvarez, *et al*., 2016; Ding, *et al*., 2018). VIPER leverages a gene expression signature and a GRN representing genomewide topological interactions between regulatory proteins and their downstream target genes, referred to as regulons. GRNs can be accurately reconstructed from gene expression data using several algorithms, e.g., ARACNe (Basso, *et al*., 2005) or SCENIC (Aibar, *et al*., 2017), among others. VIPER prioritizes regulatory proteins whose regulons are most enriched in the gene expression signature of each individual sample, by performing a gene set enrichment analysis using the analytic Rank-based Enrichment Analysis (aREA) or the novel Non-parametric analyticalRank-based Enrichment Analysis (NaRnEA) (Alvarez, *et al*., 2016; Griffin, *et al*., 2023). Moreover, the metaVIPER algorithm extends this approach to the inference of protein activity by integrative analysis of multiple, non-tissue matched GRNs, thus assuming that the transcriptional targets of each protein can be recapitulated by one or more available GRN. This is an effective strategy for the analysis of cellular populations missing a matched GRN, due to either lack of lineage information or low sample availability (Ding, *et al*., 2018). VIPER analyses have led to the identification (and validation) of several physiological or dysregulated transcriptional programs in a variety of contexts, including many cancer types and other diseases (Elyada, *et al*., 2019; Laise, *et al*., 2022; Li, *et al*., 2023; Malagola, *et al*., 2024; Mishra, *et al*., 2020; Obradovic, *et al*., 2021; Paull, *et al*., 2021; Vasciaveo, *et al*., 2023).

The recent availability of technologies to profile the transcriptome of individual cells by single-cell RNA sequencing (scRNA-seq) has provided unparallel insights into tissue heterogeneity. Critically, scRNA-seq data exhibit high sparsity and low signal-to-noise ratio due to gene dropout biases, resulting in the capture of a small fraction of the expressed genes (10-20%) in every cell (Qiu, 2020; Vlahos, *et al*., 2023). This phenomenon hinders fine grain quantitative analyses, such as the elucidation of small cell populations due to the lack of expression of few critical lineage markers. VIPER critically mitigates such gene dropout bias by robustly assessing the differential activity of regulatory and signaling proteins based on the differential expression of their downstream transcriptional targets. VIPER has been successfully applied to quantitate protein activity of >6,000 regulatory and signaling proteins in individual cells, including transcription factors and co-factors, chromatin remodeling enzymes, signaling proteins, and surface receptors (Elyada, *et al*., 2019; Li, *et al*., 2023; Obradovic, *et al*., 2021). Single-cell protein activity analysis has proven instrumental to the identification of rare cell types and novel cell states representing subpopulations that are virtually undetectable at the gene expression level, comparing favorably with flow cytometry and CITE-seq antibody-based measurements (Obradovic, *et al*., 2021). VIPER-inferred cell states have been linked to critical functional roles, including those affecting immune-evasion and progression (Elyada, *et al*., 2019; Obradovic, *et al*., 2021), disease heterogeneity and drug sensitivity in diabetes and cancer (Ding, *et al*., 2019; Li, *et al*., 2023; Son, *et al*., 2021), and stem cell biology (Malagola, *et al*., 2024). The ever-growing size of scRNA-seq datasets, due to lower reagent and sequencing cost, supports creation of extensive single-cell atlases and large-cohort analyses. Yet, they also require novel tools capable of analyzing tens to hundreds of thousands of cells. The current R-based VIPER implementations (Alvarez, *et al*., 2016; Vlahos, *et al*., 2023) is not designed to handle dataset sizes that are becoming the de facto standard for single cell studies. This is further exacerbated in their R-based NaRnEA implementation (Griffin, *et al*., 2023), which is even more computationally demanding. To allow VIPER analyses on large datasets, we introduce pyVIPER, a fast, memory-efficient, and highly scalable Python-based VIPER implementation. The pyVIPER package leverages AnnData objects (Virshup, *et al*., 2021) and is seemingly integrated with standard single cell analysis packages, such as Scanpy and others from the scverse ecosystem (Virshup, *et al*., 2023; Wolf, *et al*., 2018). Unlike previous R-based implementations, pyVIPER can directly interface with Machine Learning (ML) packages, such as scikit-learn and TensorFlow (Abadi, *et al*., 2016; Pedregosa, *et al*., 2011) to allow plug- and-play ML analyses that leverage VIPER-assessed protein activity profiles. We benchmarked pyVIPER’s performance improvements using scRNA-seq data from pancreatic ductal adenocarcinoma (Peng, *et al*., 2019) and compared to the R-based VIPER implementation (Alvarez, *et al*., 2016; Vlahos, *et al*., 2023).

## 2 Results

The package is named after the viper function, which is the Python implementation of the VIPER algorithm (Alvarez, *et al*., 2016). For each single cell, or bulk RNA-seq sample, viper estimates the activity of regulatory proteins based on the expression of their downstream targets. Inputs to viper include: 1) a gene expression matrix (rows:cells or samples, columns:genes) provided as an AnnData object or Pandas DataFrame, and 2) one or more GRN objects of a new Python class Interactome (see 2.1). The function operates in VIPER or metaVIPER mode depending on the number of GRNs provided. Users can select either one of two enrichment algorithms, aREA (Alvarez, *et al*., 2016) or NaRnEA (Griffin, *et al*., 2023), by passing the corresponding value to the ‘enrichment’ parameter (see Tutorial 1 for details). Both algorithms compute the Normalized Enrichment Score (NES) of each protein, a test statistic that represents the enrichment of the protein’s target in differentially expressed genes and serves as a proxy for the quantification of its activity (Alvarez, *et al*., 2016). In addition, NaRnEA computes the per-protein Proportional Enrichment Score (PES), a metric with values in the interval [-1; +1] that measures the effect size for each regulon’s enrichment. aREA and NaRnEA also differ in the formulation of their null models: aREA uses an empirical phenotype-specific permutation null model, while NaRnEA uses a fully analytical null model based on the maximum entropy principle (Griffin, *et al*., 2023). Compared to aREA, NaRnEA improves type I error control resulting in higher sensitivity, at the expense of increased computational time (Vlahos, *et al*., 2023). Notably, aREA and NaRnEA allow also standalone use for pathway enrichment analysis; we provide a set of functions that exploit this feature (Tutorial 2). Last, viper supports multicore parallelization to improve large dataset analysis efficiency.

### 2.1 The Interactome class

pyVIPER introduces the new Interactome class for easy GRN manipulation, as described in Tutorials 1-2. Objects of this class can be conveniently created from any Pandas DataFrame with the following columns names: *regulator, target, mor and likelihood*. The first two columns denote the genes involved in the regulatory relationship, i.e. those for which there is an edge in the network topology; mor describes the “mode of regulation”, i.e. the directionality of the relationship: a positive value means that an increase in the regulator’s activity causes an increase in target expression, a negative value means that it leads to a decrease. Last, the likelihood quantifies the weight of the association. The Interactome class provides more than 10 methods. These include: ‘filter_regulators’, to automatically filter a GRN to retain only those regulatory proteins belonging to selected ontology classes; ‘filter_target’, to focus only on specific regulatory-target interactions; ‘integrate’, to consolidate several Interactome objects into a “consensus” network that integrates multiple GRNs with similar epigenetics, such as those inferred from the same cell population in similar biological contexts but distinct datasets; ‘prune’, to prune each regulon to a specified number of targets, thus making aREA-inferred NES comparable across regulators and preventing regulators controlling more targets to numerically dominate those having fewer; ‘translate_regulators’ and ‘translate_targets’, convenient utilities to translate gene identifiers and symbols between HGNC, Ensembl genes, Entrez IDs, and orthologous genes between organisms (mouse and human). Additionally, pyVIPER offers a function ‘read_networks’ for directly importing various network outputs from widely used reverse-engineering tools, such as ARACNe-AP, ARACNe3 and SCENIC and formatting them into interactome objects.

### 2.2 pyVIPER’s modular implementation

pyVIPER has a modular architecture built around the AnnData class. Some of the main modules recall Scanpy’s framework structure and are (Fig 1A): pp. The preprocessing module embeds functions for preprocessing of gene expression and protein activity data. Among these, viper_similarity computes a matrix of NES-based cell similarity scores and allows one to specify the number of regulators for the comparison to focus on candidate MRs of the cell state; stouffer generates cluster-based protein signatures with the Stouffer’s method (Dewey, 2023); aracne3_to_regulon automatically converts a the raw output of a GRN topology reconstructed with ARACNe3 (Griffin, *et al*., 2023) into a Pandas DataFrame with regulator, target, mor and likelihood columns; rank_norm generates a gene expression signature by rank transformation of the input data (AnnData object or Pandas DataFrame) followed by median centering and mean absolute deviation-(MAD) based scaling; repr_metacells generates metacells to rescue signal lost due to gene dropouts. Metacell assembly is performed by partitioning a k-nearest neighbors (k-NN) graph into homogeneous subsets of cells and aggregating their unnormalized counts. This generates a “pseudo-bulk” profile that can be used for downstream gene expression analysis or as an input to ARACNe or other GRN-inference algorithm for datasets with median depth is <10k UMI/cell (Vlahos, *et al*., 2023). Metacell overlap is minimized by splitting all the data into N groups of neighboring cells where N is the number of desired metacells. This is followed by selecting a cell within each group for conversion to a metacell based on the minimal overlap of its k-NN with all other selected cells. (Tutorial 3). translate allows convenient utilities to translate gene names like those within the Interactome class. tl. The tools module provides functions for transformation and postprocessing of a VIPER-inferred protein activity matrix. Examples include find_top_mrs, which identifies candidate master regulator proteins from the most statistically significant proteins within each cluster; path_enr is a convenient way for users to run the VIPER algorithm to compute enrichment scores of pathways; oncomatch, which implements a tumor-to-model (e.g., organoid or cell line) fidelity analysis based on the overlap of MR proteins inferred from patient and model samples, a methodology validated in several preclinical and precision medicine studies (Vasciaveo, *et al*., 2023).

**Fig. 1.**
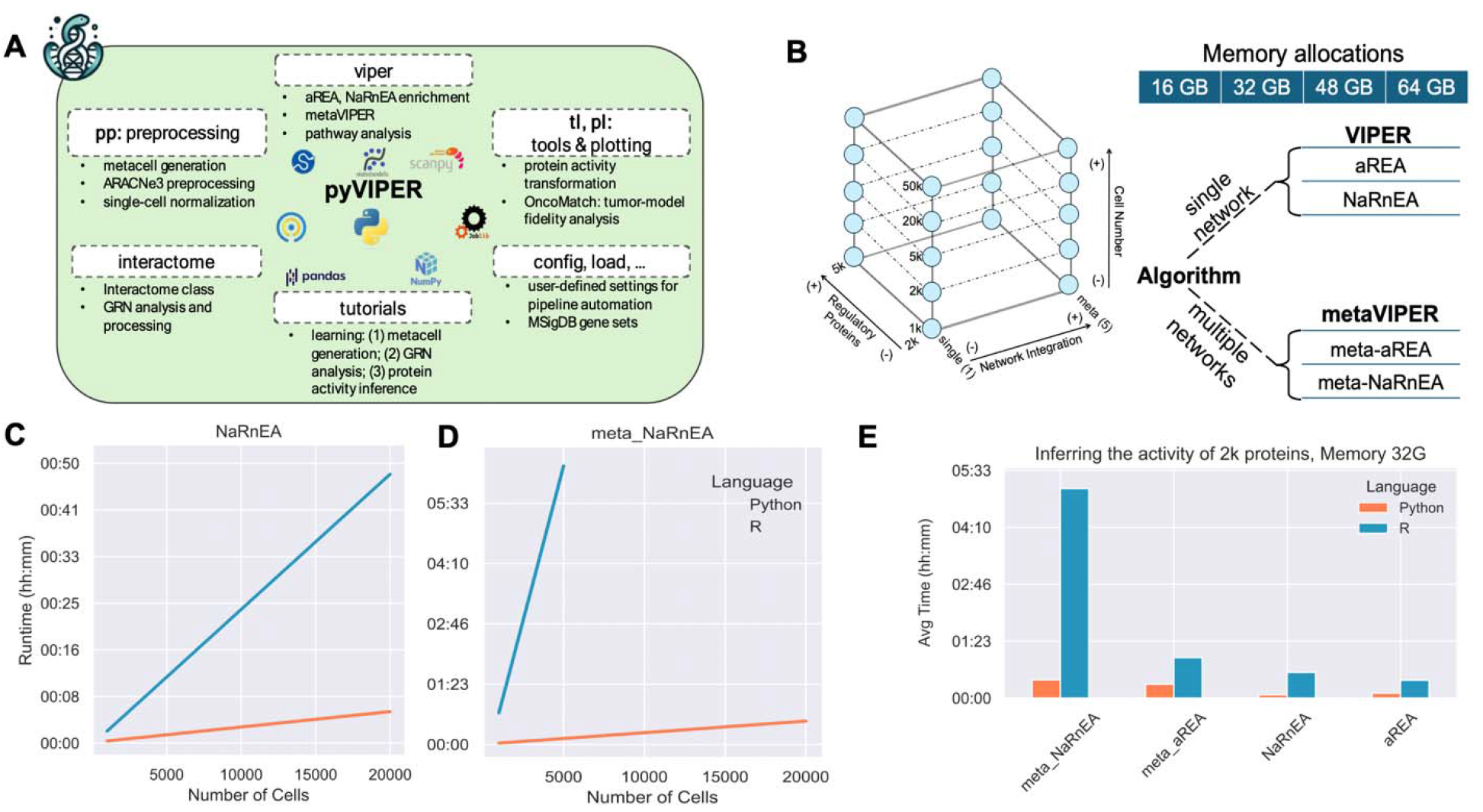
Overview of the pyVIPER package and performance comparison with R-based implementations. (A) pyVIPER modular architecture. pyVIPER includes the core viper function, several modules for Protein Activity analysis of transcriptomic data, mainly scRNA-seq, and user-friendly tutorials. (B) Python vs R runtime comparison: configurations tested. 20 configurations were generated for different numbers of i. input single-cells (1k, 2k, 5k, 20k, 50k), ii. inferred regulatory proteins (2k, 5k) and iii. input regulatory networks (1 for VIPER runs and 5 for metaVIPER runs). VIPER and metaVIPER were executed with both the aREA (meta-aREA) and NaRnEA (meta-NaRnEA) enrichment methods. Each instance was run with 4 memory allocations: 16, 32, 48 and 64 GB. (C-D) Python vs R runtime comparisons: VIPER- and metaVIPER-based Protein Activity inference of 2k regulators (NaRnEA enrichment) with 32 GB of allocated memory. pyVIPER exhibits a substantial runtime decrease compared to the R-based implementation. pyVIPER’s speedup is even more significant for the most demanding configurations (e.g., metaVIPER with NaRnEA enrichment), with runtime decrease from several hours to minutes (from 6 hours to <10 min with 5k cells). pyVIPER scales more efficiently with the number of cells, enabling the analysis of 4x cells with the same memory allocation. (E) Python vs R average runtime comparison for each enrichment algorithm: Protein Activity inference of 2k regulators with 32 GB of allocated memory shows average runtime reductions up to 15 times. All the metaVIPER runs involved 5 input regulatory networks inferred from malignant ductal, normal ductal, fibroblast, endothelial and B cell populations. Time is reported in hours, minutes (hh:mm). All tests were run on an Intel® Xeon® E5-2630 2.30GHz CPU using 1 core.

Other useful modules include pl, a wrapper for the Scanpy.pl module for visualization of gene expression, protein activity and pathway enrichment data; config, for user specifications of many default pipeline settings, such as the species being analyzed and the custom list of regulatory proteins used to automatize the analysis workflow, and load, that offers previously curated lists of regulatory and signaling proteins, and gene sets from the Molecular Signature Database (MSigDB) (Liberzon, *et al*., 2015) formatted as Interactome. To help user navigation into the numerous functionalities offered by pyVIPER, we provide 3 detailed tutorials covering relevant topics, including GRN manipulation and VIPER analysis (Tutorial 1), gene expression signature generation and metaVIPER analysis (Tutorial 2), and metacell assembly (Tutorial 3).

### 2.3 pyVIPER benchmarking performance

pyVIPER implementation offers greater scalability to large single-cell datasets comprising several hundred thousand cells, thus overcoming the memory-constraints and runtime limitations of the VIPER’s R-based implementations (Alvarez, *et al*., 2016; Vlahos, *et al*., 2023). To test the performance of the core viper function, we ran multiple benchmarks on input data of varying size (1k, 2k, 5k, 20k, and 50k cells). Each instance was generated by randomly sampling the selected number of cells from a large transcriptomic atlas of 41,986 individual cells from primary pancreatic tumors (Peng, *et al*., 2019). UMI data were normalized and transformed to Pearson residuals (Hafemeister and Satija, 2019). To mimic different biological scenarios, we computed the enrichment score of two sets of 2k and 5k regulatory proteins. Each GRNs was reverse engineered using ARACNe3. Tests were run using both aREA and NaRnEA enrichment algorithms. metaVIPER runs were run by integrating n=5 lineage-specific GRNs, each pruned to 100 targets and runtimes were recorded for several memory allocations, 16, 32, 48 and 64 GB. All the tests were run on an Intel® Xeon® E5-2630 CPU with 15M Cache, 2.30GHz, using a single core. Fig. 1B schematically represents all the configurations tested. Notably, pyVIPER demonstrated critical performance improvements compared to R-based VIPER, for all memory configurations (Fig 1E, Supplementary Fig. 1), with increasingly significant gains for the most demanding analyses (e.g. NaRnEA-based metaVIPER). As an example, metaVIPER execution on 5k cells showed a runtime decrease of up to 15 times, from approximately 6 hours to less than 10 minutes. Moreover, given the same memory allocation (32 GB), pyVIPER easily supported the analysis of 20,000 vs. 5000 for VIPER (400% improvement) (NaRnEA enrichment, Fig. 1D). Similar outcomes were observed for all memory configurations and number of inferred regulators. pyVIPER faithfully reproduced the results of the R-based implementation, as benchmarked across three single-cell datasets, with produced relative discrepancies (%) in the order of 10-14-10-15 (NES, aREA) and 10-7-10-5 (NES; PES, NaRnEA) (P value for such differences is >0.99) and protein activity profiles with Pearson’s correlation asymptotically 1 (Supplementary Fig. 2). Additional details on pyVIPER accuracy estimation are illustrated in the Supplementary Information.

## 3 Conclusions

pyVIPER addresses the increasing need for scalable and interpretable network-based analysis of large gene expression datasets, which are becoming more common in single-cell studies. pyVIPER-inferred protein activity profiles are virtually identical to those inferred with the original VIPER implementations. Yet, the computational speedup can substantially benefit researchers, particularly those who do not have access to expensive computational resources or want to perform exploratory data analyses. On a personal computer, pyVIPER can perform protein-based NaRnEA analysis of >50,000 single-cells, while enjoying a cup of tea (∼10 minutes), in contrast to the limit of 5,000 cells of R-based VIPER, making its use more accessible. The availability of new functionalities and tutorials also facilitates the systematic, highly reproducible network-based analysis of scRNA-seq data. By exploiting a modular architecture, pyVIPER is also amenable to future developments to include additional data modalities and algorithms. Moreover, pyVIPER interoperability with Scanpy and other tools from the scverse ecosystem enables easy cross-compatibility and results transfer within the Python environment. Taken together, these essential improvements suggest that the redesign and consolidation of the entire protein activity framework into the unified pyVIPER platform will increase both ease of use and deployment, providing scientists with an embedded toolkit for the network-based, systematic analysis of large-scale scRNA-seq datasets that can be further interfaced with the rapidly expanding set of Python tools for multimodal data analysis, cell-cell communication and single-cell trajectory inference.

## Data and code availability

pyVIPER code be found at https://github.com/alevax/pyviper. Documentation is available at https://alevax.github.io/pyviper/. The latest release can be installed also through PyPI for all platforms (https://pypi.org/project/viper-in-python/). Accompanying data using in the Jupyter notebook tutorials are on Zenodo (https://zenodo.org/records/10059791).

## Acknowledgements

AV and AC conceived of the study. AV supervised the study. LV implemented initial draft of the tool. AW, ZL and LZ curated all successive code upgrades, new features and benchmarks. LZ, AW, and AV wrote initial draft of the manuscript. LZ and ZL prepared the figures. LZ and AZ conceptualized and curated the tutorials. MAG, AZ and HN developed further functionalities into the Python package. All the authors edited and reviewed the manuscript. The authors gratefully acknowledge Sumit Saluja for sharing technical information on the DSBIT HPC system employed in the software benchmarking.

